# Quantitative model suggests both intrinsic and contextual features contribute to the transcript coding ability determination in cells

**DOI:** 10.1101/2021.10.30.466534

**Authors:** Yu-Jian Kang, Jing-Yi Li, Lan Ke, Shuai Jiang, De-Chang Yang, Mei Hou, Ge Gao

## Abstract

Gene transcription and protein translation are two key steps of the “*central dogma*”. It is still a major challenge to quantitatively deconvolute factors contributing to the coding ability of transcripts in mammals. Here, we propose Ribosome Calculator (RiboCalc) for quantitatively modeling the coding ability of RNAs in human genome. In addition to effectively predicting the experimentally confirmed coding abundance via sequence and transcription features with high accuracy, RiboCalc provides interpretable parameters with biological information. Large-scale analysis further revealed a number of transcripts with a variety of coding ability for distinct types of cells (i.e., context-dependent coding transcripts, CDCTs), suggesting that, contrary to conventional wisdom, a transcript’s coding ability should be modeled as a continuous spectrum with a context-dependent nature.

## Introduction

Gene transcription and protein translation are two key steps of the “*central dogma*”. While protein abundance is generally believed to be regulated by both transcriptional and translational control [1-3], it is still a major challenge to quantitatively factors contributing to transcript’s coding ability (i.e., whether a particular transcript will encode a protein and, if so, the corresponding abundance).

Benefitting from rapid development on high-throughput technology recently, several quantitative models have been proposed for modeling coding ability *in silico* based on various features in unicellular organisms [4-7]. While these models are rather accurate (e.g. the correlation of with ribosomal density has achieved 0.68[7]), heterogeneity across cells and species hinders their application in depicting translation control in mammals [8].

Multiple translation-related signatures have been reported in human and other mammal systems, revealing several gene-encoded transcription and translation regulatory features which substantially contribute to the final mRNA and protein expression levels [9-15]. Along this line, Volkova *et al*. have assessed these features and build qualitative models to discriminate coding and noncoding RNAs, as well as high- and low-translated mRNAs [16]. Trösemeier *et al*. have introduced a codon-specific translation elongation model to simulate ribosome dynamics during mRNA translation and integrate model’s parameters for protein expression prediction [17].

Here we present an experiment-backed, data-oriented computational model (named Ribosome Calculator, RiboCalc) for quantitatively predicting the coding ability (Ribo-seq expression level) of a particular human transcript (Figure 1A). Features collected for RiboCalc model are biologically related to translation control. We build the model using linear regression with Lasso penalty so that the feature parameters are easily connected to their contribution to transcript coding process. Multiple evaluations show that RiboCalc not only makes quantitatively accurate predictions but also offers insight for sequence and transcription features contributing to transcript coding ability determination, shedding lights on bridging the gap between the transcriptome and proteome. All scripts and data are available online at https://github.com/gao-lab/RiboCalc/.

**Figure 1.**
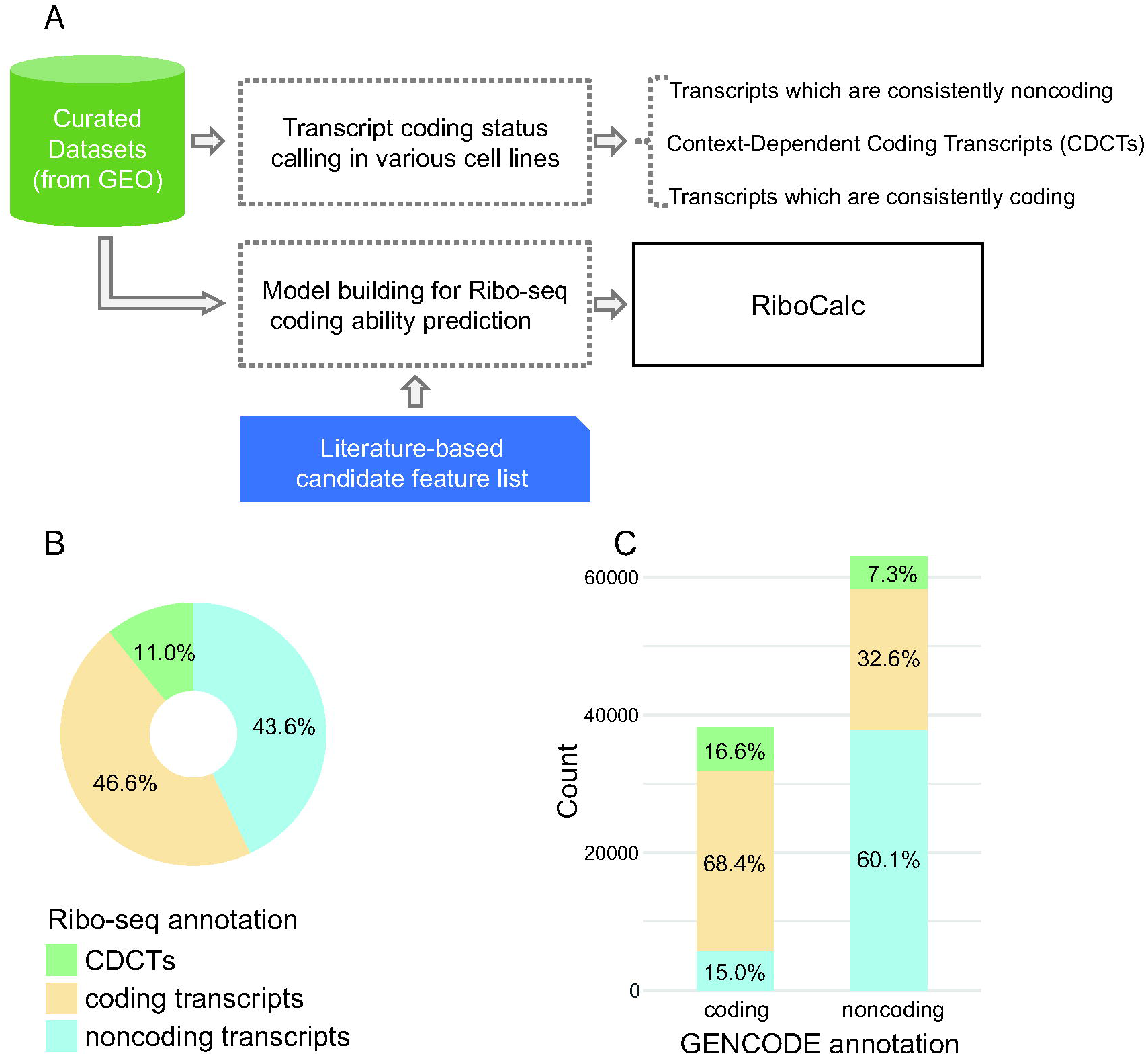
Transcript coding status classification in ribosome profiling data. **A)** Workflow of RiboCalc model building **B)** The percentage of transcripts with a particular coding ability classification. The blue fraction shows noncoding transcripts, the yellow fraction shows coding transcripts, and the green fraction shows CDCTs. **C)** Comparison between coding ability classification based on ribosome profiling with the biotypes annotated by GENCODE. The left bar shows the transcripts annotated as coding transcripts by GENCODE, while the right bar shows noncoding transcripts. The fractions of each bar correspond to the Ribo-seq based coding status calling in our analysis, and the colors of particular types are the same as those in Figure 1B.

## Materials and Methods

### Ribosome profiling data collection

We retrieved human data published since 2012 from RPFdb[18] and the NCBI BioProject database[19] by searching for the terms “ribosome profiling”, “ribosome profile”, “Ribo-seq”, “ribosome footprint” and “RPF”. We then manually selected Ribo-seq samples with paired RNA-seq data and without treatment interfering with translation. As a result, 61 datasets were retained from 30 studies covering 22 different human tissues or cell lines (Supplementary Table 1). The pipelines of transcriptome analysis for RNA-seq and Ribo-seq data are at https://github.com/gao-lab/RiboCalc/blob/master/feature_calculation/RNAandRibo-seq_processing.txt.

### Mass spectrometry (MS) data analysis

To build a reliable coding ability prediction model, the first step is to identify *bona fide* coding transcripts, especially given several recent reports that a lot of annotated noncoding RNAs (ncRNAs) were found to encode peptides [20-22]. We selected a Ribo-seq based coding gene identification method which covered more than 90% of MS-based callings (Supplementary Figure 1, Supplementary Table 2 and 3).

To find the criteria for coding gene identification covering most MS observations, we compared the ribosome profiling based results with the MS results. As reported, mass spectrometry identifies proteins with high specificity but limited sensitivity[23]. Therefore, taken MS results as golden positive calls, we selected criteria to ensure that Ribo-based results covered 90% MS calls while with less false positives. The MS dataset for assessment was PXD002395 from the PRIDE database [24]. The overlapping cell lines in this MS project and our ribosome profiling data were HEK 293, HeLa and U-2 OS cells, so only MS data from these 3 cell lines were analyzed (Supplementary Table 2).

We then compiled a human protein database by adding theoretically translated peptides of transcript putative ORFs and protein-coding transcript translation sequences annotated by GENCODE release 24. Redundant protein sequences with identity higher than 90% were trimmed using CD-hit[25] in the database. We used pFind3 as the search engine [26]. 286 common contaminant proteins were automatically added into the original protein database by pFind. The reverse protein sequences were used as a decoy database for false discovery rate (FDR) control. The search parameters were as follows: 1) Trypsin/P digestion. 2) The precursor tolerance and fragment tolerance were set to 20 ppm and 20 ppm, respectively. 3) The search included variable modifications of methionine oxidation and N-terminal acetylation. 4) Minimal peptide length was set to six amino acids and a maximum of two missed cleavages was allowed. Peptides were filtered with a FDR threshold of 1%. We identified a gene as coding in MS when supported by at least one protein with a pFind Q-value less than 0.01 and more than 5 peptide fragments.

The approaches that we used for translated ORF scanning in ribosome profiling data were RiboCode[27] and ribORF[28]. Genes with translated ORF were identified as coding in Ribo-seq. We compared the MS and ribosome profiling results in corresponding cell lines (Supplementary Table 3). MS-based coding genes were taken as positive calls. Abbreviations in the equations below are as follows: FN, false negative; FP, false positive; TN, true negative; and TP, true positive.

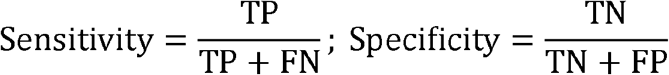

RiboCode showed a lack of consistency with MS analysis (Supplementary Table 3), while ribORF, with a p-value cutoff of approximately 0.5, could achieve a sensitivity of 90% (Supplementary Figure 1). Thus, we adopted ribORF with a p-value higher than 0.5 as an approach for translated ORF identification.

Transcript’s coding ability contributes to the protein abundance determination process significantly. During the last two decades, the most common method for large-scale experimental determination of transcripts’ coding ability is to measure the abundance of the corresponding proteins based on MS methods. Nevertheless, MS-based protein identification is less sensitive to short peptides and peptides expressed at low levels[29]. In addition, MS analyses often adopt a database-dependent search strategy that ignores genomic mutations and RNA editing events, hampering identification of the complete protein pool[30]. Meanwhile, by sequencing the RNA fragments protected by ribosomes, Ribo-seq measures translational activity in a quantitative manner with base resolution. Hence, Ribo-seq derived transcript density is an appropriate measure for evaluation and estimation of coding ability. During ribosome profiling data processing, we used MS results as the gold standard for determining the threshold of Ribo-seq based methods. This strategy ensured reliable, sensitive and precise identification of MS-supported protein-coding genes in the Ribo-seq data, providing an accurate training set for model building.

### Identification of translated ORFs

After scanning translated ORF based on ribosome profiling data, we made pairwise comparisons between overlapping ORFs to select a most likely translated one. Translated ORFs were identified according to the following procedure (Supplementary Figure 2): Scenario 1) overlapping translated ORFs with unique regions covered by ribo-reads were both retained. Scenario 2) the ORFs without unique ribo-reads covering region were filtered out when their overlapping ORFs had. Scenario 3) if the unique regions of both overlapping ORFs were not covered by ribo-reads, the shorter ORF was left. Thus, we compiled a non-redundant catalog of translated ORFs. Since the translated ORFs defined by us were based on ribosome profiling data, they might differ from the main ORFs in canonical annotation (Supplementary Figure 3). If there were no ribosome evidences (such as independent testing without Ribo-seq data), the longest ORF was taken as the putative translated ORF.

### Calling transcript’s coding status

As we scanned translated ORFs in Ribo-seq, we identified coding transcripts (have translated ORF) for each sample. Then, a transcript would be called as “noncoding” in a particular sample only if 1) not covered by any ribo-reads, 2) its expression abundance is higher than the lower bound of called “coding” ones in this sample. The expression threshold was set to the 300th quantile of transcripts per million (TPM) for all coding transcripts in the corresponding sample, so, at this abundance, translated ORFs could be effectively detected. By using the 300th quantile as a threshold, a large number of expressed transcripts (96,968) were removed (Supplementary Figure 4), indicating that the process is stringent enough for ncRNA identification. Another circumstance is that a transcript is covered by Ribo-seq reads but no translated ORF is identified. We also removed this kind of transcripts (1,031) because of lacking reliable evidence for coding.

As described above, we identified coding and noncoding transcripts in each sample. Transcripts with translated ORF in every expressed sample were classified as coding, while the transcripts consistently without translation were classified as noncoding. In addition, several transcripts showed coding in some samples but were noncoding in other samples. We defined these transcripts as CDCTs (context-dependent coding transcripts). In further model building, we retained genes with all isoforms as coding and selected one representative isoform (expressed in the most samples) for the gene into the training data (Supplementary Figure 5).

### Collection of classified features based on biological knowledge

Based on biological knowledge, the candidate features were collected from manual literature survey. Our aim is to depict translation control *in vivo*, so the first criterion for feature collection is able to be explained by translation-related biological process. The second criterion is the feature value should be easily encoded by sequence and transcription information. As a result, we collected 221 features and grouped them into 5 translation-related processes. 1) Expression abundance: in addition to RNA-seq expression level, miRNA targeting also affect RNA abundance. Thus, we scanned miRNA target sites on 3’UTR and incorporated it into a feature. 2) Translation initiation: the RNA folding energy and sequence context around the translation initiation site are related to the transcripts’ coding ability, so we used these features in RiboCalc. 3) Translation elongation: the translation elongation rate is often altered by the adaption between codon usage and the corresponding tRNA abundance[10]. Therefore, the frequency of 64 codons, as well as some other indexes describing codon usage bias, were used in this class of features. 4) Translation regulators: the abundance of translation regulatory factors are related to translation level of RNAs[31], thus, expression level of translation-related genes annotated by GO were taken into account as features. 5) Transcript structure: other sequence features are also related to transcript coding ability, such as length and UTR GC content. The mechanism of these features might not be clearly validated with experiments but we added them into the list.

Considering the genomic mutations and RNA editing, we modified transcript reference sequences with variations called by GATK[32] in RNA-seq, and calculated feature value based on modified sequences. The mutations causing start codon loss were ignored (see “Mutated transcript sequence identification with RNA-seq data” at https://github.com/gao-lab/RiboCalc/blob/master/feature_calculation/RNAandRibo-seq_processing.txt). The ORFs that we used for feature calculation are described in Supplementary Figure 3. All the features and the calculation method are shown in Supplementary Table 4. We provide feature calculation scripts at https://github.com/gao-lab/RiboCalc/tree/master/feature_calculation.

### Building cell-specific models

We chose 5 cell lines for cell-specific model building. The 5 cell lines were the most common ones among our collected resources (Supplementary Table 1) and could represent 5 different tissues. We used Ribo-based coding transcripts for model building and removed redundant sequences with more than 90% identity using CD-hit[33]. We randomly selected 3,000 transcripts as training data and the rest as testing data for each model. The models with selected features were built through linear regression with the Lasso penalty. The feature data and model building script are at https://github.com/gao-lab/RiboCalc/tree/master/cell_specific_model.

### RiboCalc model building

By adding *trans-* features of translation regulators, we built an “environment-aware” model for quantitative prediction of coding ability globally and named the model RiboCalc. The detailed methods of model building were as follows:

We pooled coding transcripts from all cell lines together. If identical transcripts expressed in more than one ribosome profiling sample, we selected the one with median TPM in the corresponding RNA-seq data. The dataset consists of 8,193 transcripts, and we randomly split them into 5,000 training cases and 3,193 testing cases.

All the values of each feature were scaled to the interval of [0, 1] for training data as following equation (min-max normalization).

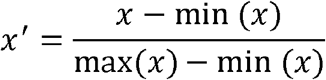

Since gene-level expression abundance estimation was reported to be more accurate than isoform level [34-36], we took gene TPM for calculation. Since different studies had a various distribution of expression level in RNA-seq and Ribo-seq, to pool all the data together, we adopted cross-sample normalization for RNA-TPM and Ribo-TPM. The RNA-TPM and Ribo-TPM were normalized with TPM of the housekeeping gene *HPRT1* using the following equation. *HPRT1* was selected based on the work of Valente *et al*.[37].

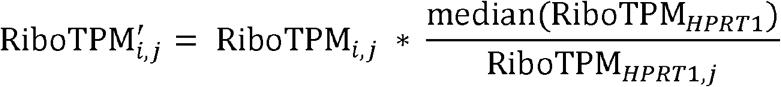

The Ribo-TPM of transcript *i* in sample *j* was scaled by the ratio of the *HPRT1* Ribo-TPM in sample *j* with the median *HPRT1* Ribo-TPM among all the samples. The same normalization strategy was also applied to RNA-TPM.

We first removed highly correlated features (Pearson’s *r* above 0.9) with “findCorrelation” in the R caret package[38]. Then, feature selection was implemented through a linear model with Lasso regularization. We searched the parameter λ with the minimum mean squared error (MSE) in 5-fold cross validation. The Lasso regression was implemented by the glmnet package[39] in R. We provide raw data and script at https://github.com/gao-lab/RiboCalc/tree/master/RiboCalc.

### Human model comparison

#### OCTOPOS [17]

We download the raw data OCTOPOS used for HEK 293. The RNA-seq data was from https://www.ncbi.nlm.nih.gov/geo/query/acc.cgi?acc=GSE38356. Protein abundance data was downloaded from https://pax-db.org/dataset/9606/329/. Since correlation calculation would be affected by data size, we randomly selected same amount of transcripts (461) with OCTOPOS as testing set. We provide the script of feature calculation and model testing from OCTOPOS data at https://github.com/gao-lab/RiboCalc/blob/master/feature_calculation/script/test_OCTOPOS.sh. Users could follow the script to apply RiboCalc on their own data.

#### Li’s human model [40]

We downloaded their data “Additional file 2” at https://doi.org/10.1186/s13059-019-1761-9. The UTR and CDS sequences were obtained from Ensembl Genes 104 with the transcript IDs. Since Li’s human model used RPKM as expression level which was hardly transformed into TPM without knowing the full transcriptome[41], we retrained RiboCalc with their data. We randomly selected 2,000 transcripts as testing data and the rest as training data. The generated feature data and testing script are at https://github.com/gao-lab/RiboCalc/tree/master/human_model_comparison/LiJJ.

#### Sample’s model [42]

We downloaded the model from https://github.com/pjsample/human_5utr_modeling/tree/master/modeling/saved_models/main_MRL_model.hdf5. Given that Sample *et al*.’s model requires the input sequence length to be 50, we generated 50nt fragments (window size = 50, step size = 1) of 5’UTRs of RiboCalc testing data. The 5’UTR sequences were downloaded from Ensembl Genes 104 and the transcripts without 5’UTR annotation were removed in the testing set. We used the average predicted value of all 50nt windows from 5’UTR sequences as the final predicted value of the transcripts. See https://github.com/gao-lab/RiboCalc/tree/master/human_model_comparison/SamplePJ.

### Comparison with Li’s model in yeast

Li *et al*. used transcript sequence features to predict translation rate (TR) in yeast [6]. In their definition, TR is defined as the number of protein molecules translated per mRNA molecule, which is the ratio of ribosome density (also abbreviated to Ribo-TPM for uniformity with RiboCalc) to RNA expression abundance in Ribo-seq. Thus, RiboCalc could predict TR by dividing RNA abundance into the output as the equation below.

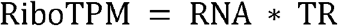

The yeast transcript sequences, ORFs, expression abundance and ribosome density were obtained from the “nar-00812-a-2017-File019.csv” file in the supplementary data downloaded from https://doi.org/10.1093/nar/gkx898. Since the RiboCalc prediction depends on the 3’UTR sequence which is not provided in Li’s data, we fetched the 3’UTR sequences from the Saccharomyces Genome Database (SGD) [43] and removed 909 transcripts without 3’UTR annotation.

To build the RiboCalc yeast model, we randomly split the remaining 1,541 transcripts into 1,000 training cases and 541 testing cases. After considering the systematic differences between yeast and human models, we removed several human-specific features from the yeast model (Supplementary Table 5). The yeast model training approach was identical to the RiboCalc human model. To make a fair comparison, we retrained Li’s yeast model using the new training set by strictly following their description. The performance of Li’s retrained model was similar to their original report (R^2^ of TR are both 0.80). The correlations in Table 3 were calculated from the testing set. See https://github.com/gao-lab/RiboCalc/tree/master/RiboCalc_yeast for raw data and script.

### Ribo-lncRNA analysis

Long noncoding RNAs (lncRNAs) associated with ribosomes are abbreviated to ribo-lncRNAs here. The ribo-lncRNAs from Ruiz-Orera *et al*. were identified from the GSE22004 dataset in the NCBI GEO database [20]. Thus, we used GSM546926 as the Ribo-seq sample and GSM546927 as the RNA-seq sample in GSE22004 for RiboCalc analysis. The “top coding score lncRNAs”, “lncRNAs with homologies” and “young codRNAs” were identified by Ruiz-Orera *et al*. and downloaded from https://www.ncbi.nlm.nih.gov/pmc/articles/PMC4359382/bin/elife03523s002.xls. The “non-ribo-lincRNAs” and “ribo-lincRNAs” and “other codRNAs” were from our analysis of the data. “Non-ribo-lincRNAs” refers to long intergenic noncoding RNAs (lincRNAs) without ribo-reads, while “Ribo-lincRNAs” are the lincRNAs covered by ribo-reads. The “other codRNAs” are protein-coding transcripts with translated ORFs in ribosome profiling excluding the young codRNAs from the original report. In this study, transcripts with FPKM lower than 0.2 were excluded.

For ribo-lncRNAs identified by Zeng *et al*. [44], their resources were included in the Ribo-seq samples collected by us, except for the data associated with drug treatment. Therefore, we directly retrieved the features of those ribo-lncRNAs in our data. The four classes of ribo-lncRNAs in Figure 3C were downloaded from https://www.ncbi.nlm.nih.gov/pmc/articles/PMC5975437/bin/12864_2018_4765_MOESM10_ESM.xlsx. See https://github.com/gao-lab/RiboCalc/tree/master/ribo-lncRNA for raw data and script.

## Results

### Transcripts’ coding status identification in ribosome profiling data

To accurately quantify the coding ability of transcripts in various cells, we retrieved 61 pairs of reliable Ribo-seq data coupled with RNA-seq data from the NCBI GEO database [19], covering 1 tissue and 21 cell lines (Supplementary Table 1). The expression abundance in the corresponding Ribo-seq data (abbreviated to Ribo-TPM) were employed as the quantitative metric for the coding ability. For accurate prediction of coding ability, the first step is to obtain protein-coding transcripts for model building. Thus, by applying rigorous filtering criteria, we called translation status for 101,170 out of 199,169 GENCODE gene models (the rest were filtered out due to either a lack of expression signals in the chosen samples (96,968) or a failure of calling translated ORF reliably (1,031), see Supplementary Figure 5 for detailed procedure). Among the 101,170 transcripts, the translation status of 46% were found to be “coding” while 43% were “noncoding” in all samples (Figure 1B). Interestingly, we also found that 11% of the transcripts exhibited diverse coding ability among cell lines (i.e., coding in some cell lines but noncoding in others), and we named them context-dependent coding transcripts (CDCTs).

### RiboCalc: predicting transcript coding ability in human

To identify features contributing to coding ability determination, we first compiled a candidate list of intrinsic (“*cis*-”) and contextual (“*trans*-”) features after systematic literature survey (Supplementary Table 4, also see “**Collection of classified features based on biological knowledge**” in “**Materials and Methods**”). The intrinsic features represent transcript sequence characteristics. And we grouped them into three categories: “translation initiation”[11], “translation elongation”[45] and “transcript structure” [6, 13] based on the underlying biological process. The contextual features were collected to depict the environment for translation in the cell and all belong to the category “expression abundance” [46]). We then incorporated these features together to build cell-specific models for five representative cell lines with a Lasso regression based feature selection. All models works well (Table 1 and Supplementary Figure 6), highlighting the effectiveness of these selected features (Supplementary Table 4).

**Table 1.** Ribo-seq resources and transcript dataset size for cell-specific model building.

| Sample ID | Cell line | Bioresources | Data size | Pearson's $r$ with Ribo-TPM |
| --- | --- | --- | --- | --- |
| SRR1803151 | GM12891 | B lymphocyte | 4,781 | 0.785 |
| SRX870805 | HEK 293 | Embryonic kidney cell | 6,057 | 0.836 |
| SRR970565 | HeLa | Cervical cancer cell | 6,440 | 0.886 |
| SRR627625 | BJ | Foreskin fibroblast | 5,746 | 0.830 |
| SRR3208870 | hESC.2 | Embryonic stem cell | 6,260 | 0.862 |

We further tried to incorporate the effect of cell “environment” by introducing expression level of *trans-*factors (i.e., translation regulators) in the corresponding sample (“Translation regulators” in Supplementary Table 4). The “environment-aware” model (RiboCalc) accurately predicts coding ability (Ribo-TPM) globally, with *r* = 0.81 in in testing data from all 22 cell lines (Figure 2A). The performance of RiboCalc is comparable with 5 cell-specific models (Supplementary Figure 6), suggesting its prediction efficiency across cells by adding “environment” features.

**Figure 2.**
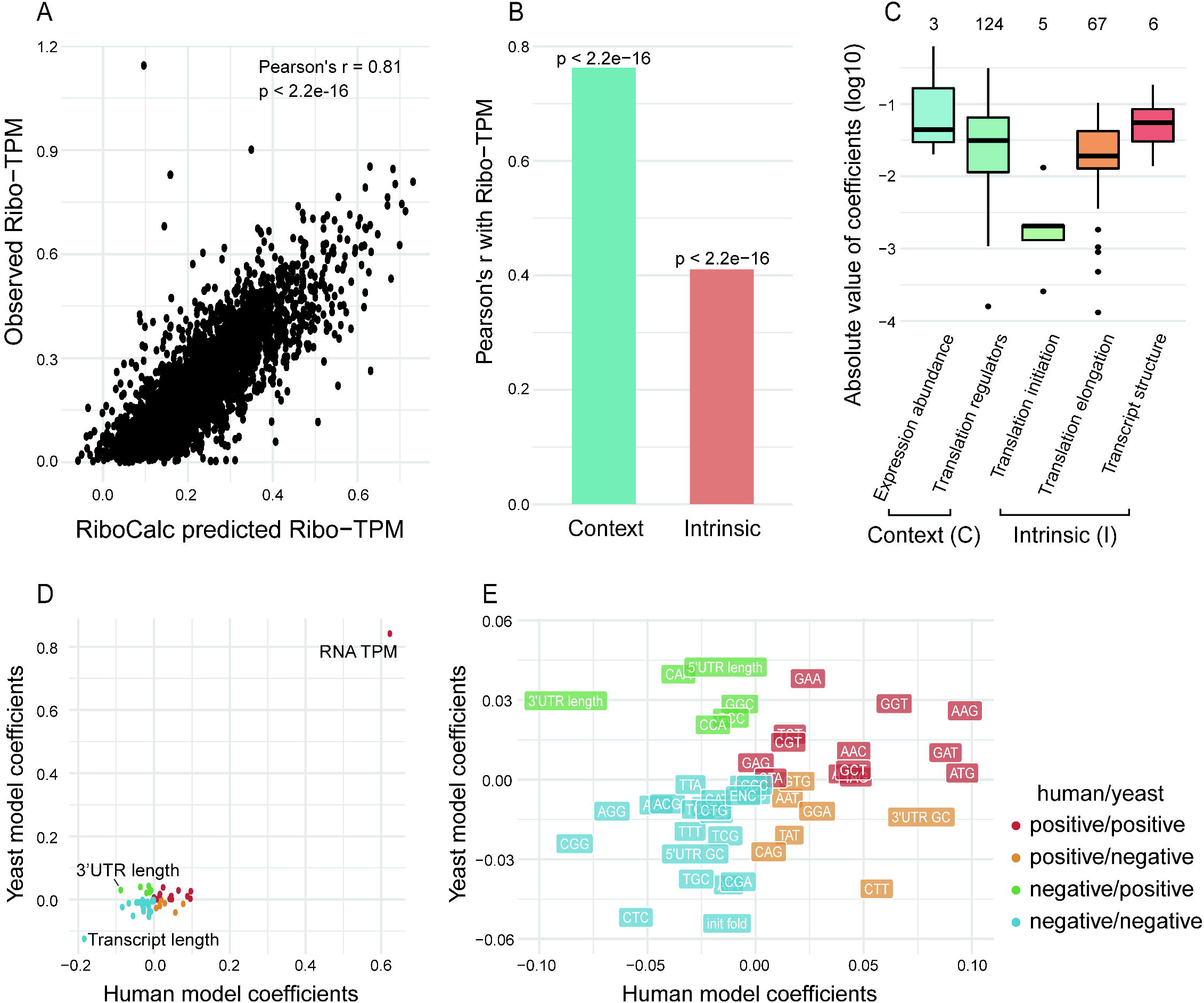
Model performance and feature contribution of RiboCalc. **A)** Scatter plot of RiboCalc predicted and observed values. The x axis shows the observed Ribo-TPM of coding transcripts in the testing set, while the y axis shows the corresponding Ribo-TPM predicted by RiboCalc. Pearson’s *r* and the significance level were calculated between x and y scores. **B)** Effectiveness of intrinsic and contextual features in RiboCalc. The bars show Pearson’s *r* between the Ribo-TPM predicted by each single class of features and the observed value. The p-value above the bars indicates the significance of the correlation. **C)** Feature importance of 5 processes in RiboCalc. The boxes show the feature coefficient distributions of the 5 translation-related processes. Since all the feature values were scaled to [0, 1], the feature coefficients could be taken as contributions to coding ability prediction in this plot. The numbers above the boxes are the corresponding feature numbers of each class. **D)** Scatter plot of feature coefficients in RiboCalc human and yeast models. The x axis shows the feature coefficients in the RiboCalc human model. The y axis shows the corresponding feature coefficients in the RiboCalc yeast model. The colors of the points represent the four quadrants. The black text labels the feature points of transcript length, 3’UTR length and RNA TPM. **E)** The feature names and coefficient values of RiboCalc human and yeast models in each quadrant. The text on the labels stand for the feature names (Supplementary Table 4). The codon name stands for codon usage frequency, “init_fold” means translation initiation sequences’ minimum free energy (MFE) predicted by RNAfold. The points of transcript length and RNA TPM are not shown in this plot due to the limitations of the axes. The x axis, y axis and colors are the same as those in Figure 2D.

We compared RiboCalc’s performance with Sample *et al*.’s model [42] and Li *et al*.’s model [40] in human. Li’s human model predicted TR (translation rate) which could be calculated as the ratio between Ribo-seq and RNA-seq abundance. RiboCalc predicted TR with Pearson’s *r* = 0.66, which is higher than Li’s human model (*r* = 0.64, Supplementary Table 6). Sample et al. built a deep learning model to predict ribosome loading. Since Sample *et al*. only provided model data of 50nt 5’UTR sequences which are not sufficient for RiboCalc prediction, we applied Sample’s model directly on RiboCalc testing data. It showed a much lower correlation (*r* = 0.18, Supplementary Figure 7) than RiboCalc (*r* = 0.81, Figure 2A). Therefore, RiboCalc accurately predicts ribosome density with intrinsic (“*cis*-”) and contextual (“*trans*-”) features.

### Parameters of RiboCalc are interpretable by biological impact on translation

As RiboCalc is a linear model fitted by normalized feature values, features’ coefficients quantify their contribution (e.g. positive coefficients suggest facilitation of coding ability, while negative coefficients suggest an adverse effect). A manually checking in literatures effectively connects model parameters with prior biological knowledge (Table 2). For instance, the feature with the highest positive coefficient is the RNA abundance (i.e. TPM), confirming existing reports on the dominant influence of transcript expression level on protein translation [46, 47]. Similarly, being consistent with the observation that longer transcripts reduced the number of dropped ribosomes diffusing to the translation initiation site as well as mRNA circularization[13], the model also demonstrates significant adverse effect for the length of transcript.

**Table 2.**
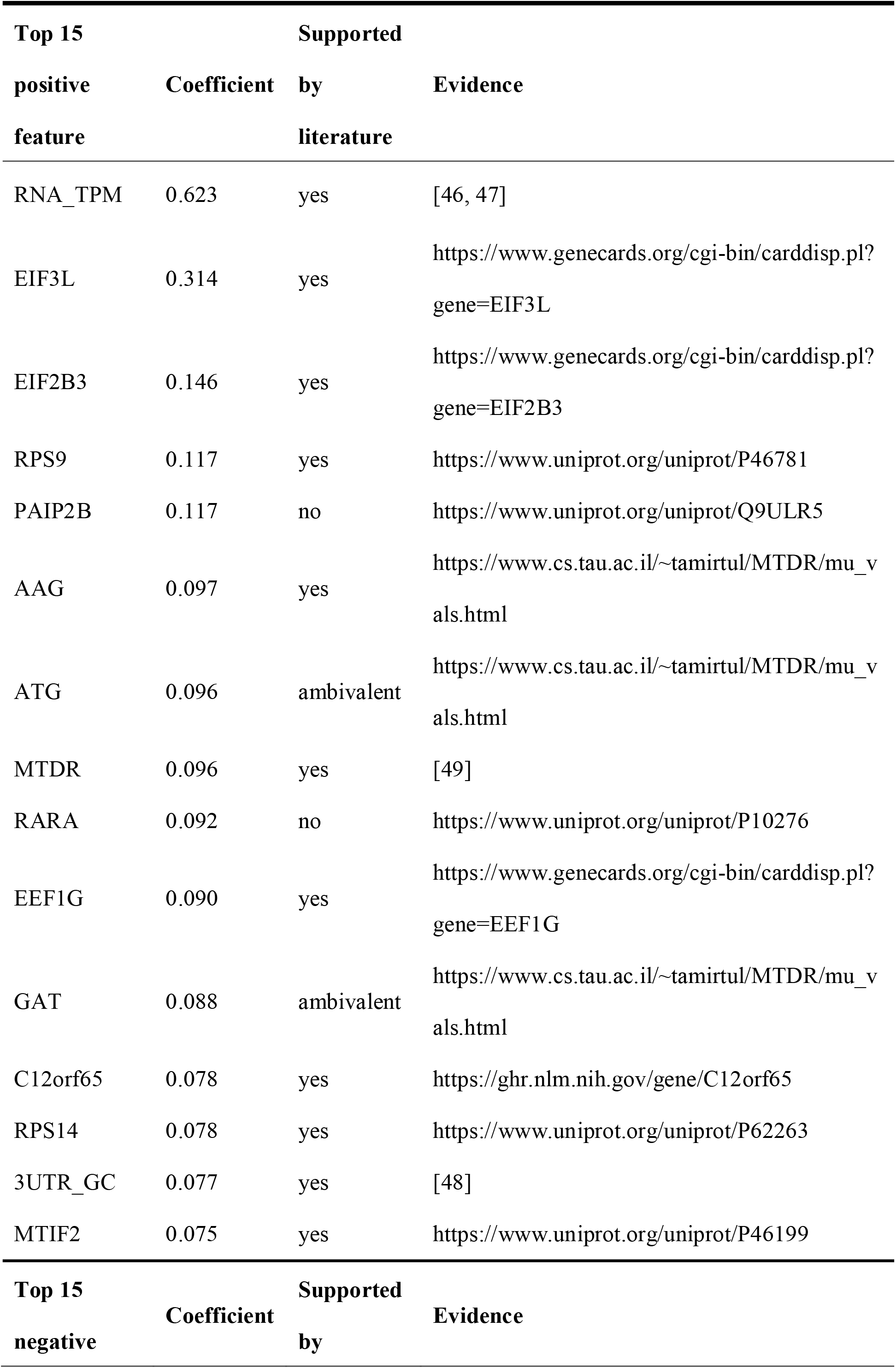
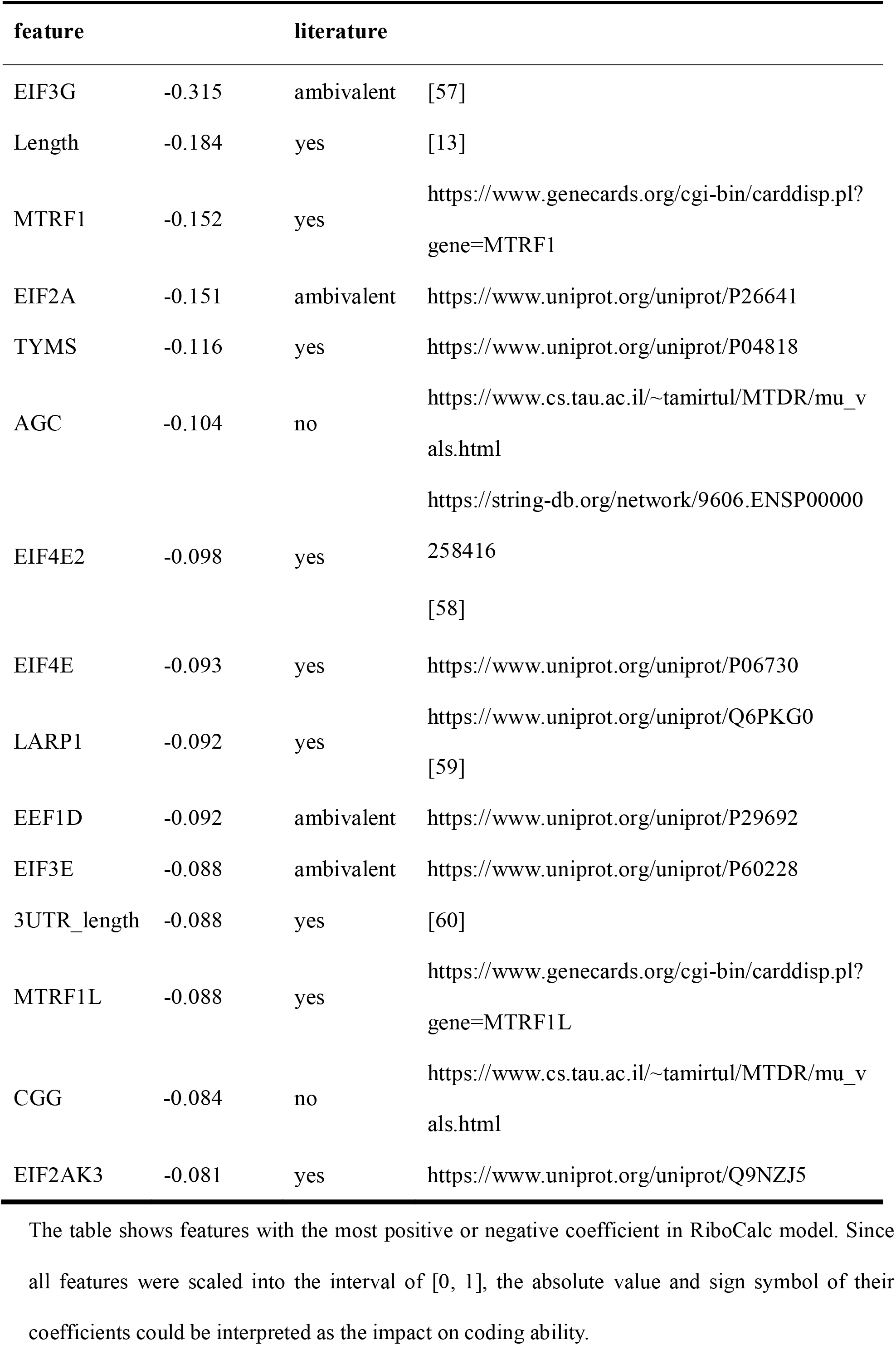
Feature coefficient with greatest contribution in RiboCalc and the effect on translation in the literature.

Based on biological knowledge, we grouped all features into 5 translation-related processes, as translation initiation, translation elongation and transcript structure for intrinsic features from sequence, and expression abundance and translation regulators as contextual features for environment. By calculating Ribo-TPM through a single class of features with RiboCalc, we compared their performance with the correlation between predicted and observed value. The predicted results of both intrinsic and contextual features showed a significant correlation with the observed Ribo-TPM (Figure 2B). And consistent with previous studies [46], expression abundance showed the greatest importance for coding ability in RiboCalc (Figure 2C).

To further validate the model’s effectiveness, we applied it to unicellular organism yeast based on published dataset[6], and found that the original RiboCalc model accurately predicted both coding ability and translation rate, with comparable performance to the state-of-arts model in yeast (Table 3, see “**Comparison with Li’s model in yeast**” section in “**Materials and Methods**” for details). Given the systematic differences between human and yeast, we also retrained a RiboCalc yeast model, with yeast data as input but adopting exactly the same feature set and fitting procedure as in the human model. As expected, the RiboCalc yeast model further improved performance overall (Table 3).

**Table 3.** Correlation between predicted and observed values in RiboCalc human, RiboCalc yeast and Li’s model.

| Predicted value | Model | Pearson's $r$ | Spearman's $r$ |
| --- | --- | --- | --- |
| TR | RiboCalc human | 0.759 | 0.762 |
|  | RiboCalc yeast | 0.886 | 0.898 |
|  | Li yeast | 0.874 | 0.887 |
| Ribo-TPM | RiboCalc human | 0.974 | 0.969 |
|  | RiboCalc yeast | 0.987 | 0.984 |
|  | Li yeast | 0.986 | 0.982 |

Intriguingly, we found that, through the same set of model features adopted, multiple coefficients of the RiboCalc yeast model differ from those of the human model. The feature coefficient with the largest difference is the length of 3’UTR (Figure 2D). Recently, Fu *et al*. demonstrated that a longer 3’UTR increased the possibility of miRNA targeting in mammalian cells, resulting in a reduction in protein translation[12], whereas no clear evidence for the pervasive existence of miRNAs in yeast. Consistently, the coefficient for the 3’UTR length in the human model is a large negative value and close to zero in yeast (Supplementary Table 7, also see Figure 2D and 2E for another case on codon usage). We believe that divergent pattern reflected by RiboCalc models could facilitate investigating the inter-species discrepancy of translation regulatory mechanisms.

## Discussion

To model transcript coding ability quantitatively, we proposed and implemented RiboCalc. RiboCalc not only effectively predicts coding ability but also reveals several intriguing novel hints for deconvoluting the translation control mechanism. For example, according to the model, the GC content at 3’ UTR presents a positive contribution to the coding ability. Coupling with recent report on AU-rich 3’UTRs leads to decreased stability of RNAs[48], we could reasonably infer that the 3’UTR with higher GC content would promote translation efficiency by slowing RNA decay. Meanwhile, we also notice a few inconsistencies between model-estimated coefficients and existing literature. For example, a reported translation-suppressing gene, *PAIP2B*, showed a positive contribution to coding ability in our model, with statistically significant positive correlation detected for its abundance and the median Ribo-TPM of translatable genes in the corresponding sample (Supplementary Figure 8, also see Supplementary Figure 9). All these observations shed lights on further mechanism study and validation.

These insights could lead to several potential applications like improving existing codon optimization tool [17, 49]. A direct comparison with OCTOPOS, a recent published mechanism-oriented codon optimization tool [17], found that the RiboCalc model could effectively predict protein abundance (*r* = 0.63, HEK293 human dataset, vs. *r* = 0.61 reported in the original paper, Supplementary Figure 10A), even given the fact that the RiboCalc model was trained to predict Ribo-seq level, instead of protein abundance measured in MS as what the OCTOPOS did. Of interest, when being retrained with OCTOPOS HEK293 dataset, the RiboCalc model showed improved prediction accuracy (*r* = 0.73, Supplementary Figure 10B, also see “**Human model comparison**” in “**Materials and Methods**”), highlighting RiboCalc’s potential to pinpoint novel translation-regulation-related features.

The RiboCalc model suggests that both intrinsic (“*cis*-”, like the transcript sequence) and contextual (“*trans*-”, like expression level of transcript and translation regulators) features contribute to the transcript coding ability decision in cells. Intriguingly, we found a number of transcripts exhibited diverse coding ability among cell lines (CDCTs, Supplementary Figure 11, also see detailed list at https://github.com/gao-lab/RiboCalc/blob/master/CDCT/CDCT_transcript_list.tab). Among the protein-coding transcripts annotated by GENCODE, 15% were classified as noncoding and 17% as CDCTs; meanwhile, one-third of GENCODE-annotated ncRNAs were classified as coding, and 7% as CDCTs (Figure 1C, also see the list at https://github.com/gao-lab/RiboCalc/blob/master/CDCT/CDCT_lincRNA_TransLnc_evidence_number.tab). The model shows that CDCTs under coding context get higher RiboCalc scores than these under noncoding context (Figure 3A). Consistently, canonical protein coding genes have been validated experimentally that protein expression ability could be varied or even silenced without altering mRNA transcribing [50-52] and several annotated lncRNAs were found to be associated with ribosomes (abbreviated to ribo-lncRNAs) [20, 44, 53] and encode functional peptides [54-56]. RiboCalc model confirms that reported ribo-lncRNAs as well as lncRNAs with high homologies and top coding score[20, 44] have coding ability significantly higher than that of non-ribo-lncRNAs, and close to those of young experimentally validated coding RNAs (Figure 3B and 3C, also see Supplementary Figure 12 for more analysis on GENCODE transcripts). Collectively, these results suggest that transcript’s coding ability should be modeled as a context-dependent continuous value, rather than a certain binary class.

**Figure 3.**
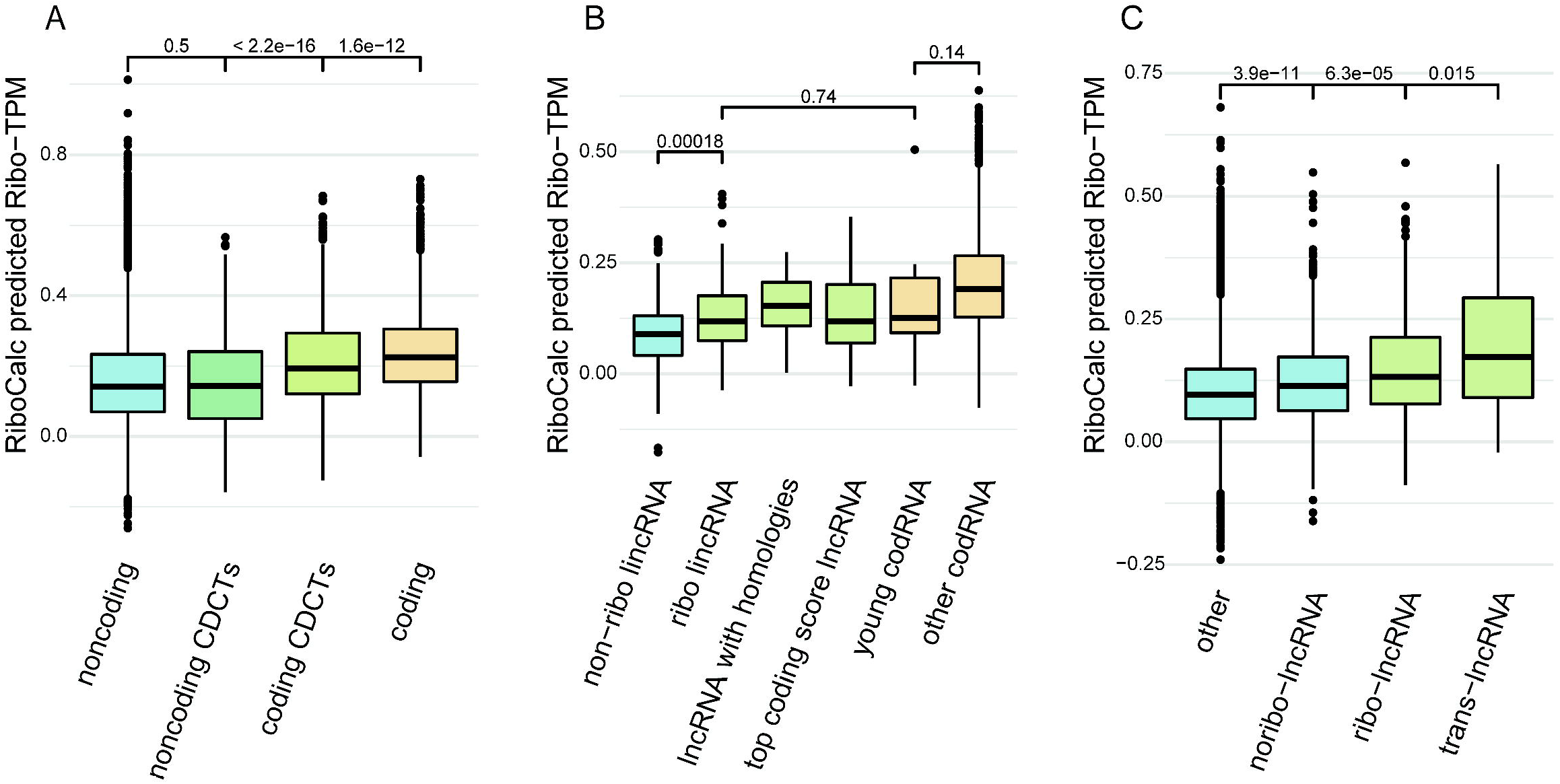
Biological interpretation of transcripts with ambiguous coding ability from the RiboCalc model. **A)** RiboCalc prediction of transcripts with a particular coding ability classification. “Noncoding” refers to the noncoding transcripts identified in Ribo-seq, “coding” is the testing data of RiboCalc. The “coding CDCTs” are CDCTs present under coding context (observed as coding in particular samples), and “noncoding CDCTs” are under noncoding context. **B)** Prediction of ribo-lncRNAs from Ruiz-Orera *et al*. in RiboCalc. The boxes show the predicted Ribo-TPM distribution of a particular class of RNAs. The “non-ribo-lncRNA” are lincRNAs without ribosome coverage, while “ribo-lncRNA” are covered by ribo-reads. The “top coding score lncRNA” are lncRNAs with the highest sequence similarity with protein-coding transcripts. The “lncRNA with homologies” are lncRNAs conserved among species. “Young codRNA” are validated coding RNAs with a short evolutionary history, while “other codRNA” are the rest coding RNAs. **C)** Prediction of ribo-lncRNAs from Zeng *et al*. in RiboCalc. The boxes show the predicted Ribo-TPM distribution of a particular class of RNAs. The “trans-lncRNA” are translated lncRNAs, “ribo-lncRNA” are lncRNAs only covered by ribo-reads, “non-ribo-lncRNA” are lncRNAs without ribo-reads, and “other” refer to unexpressed lncRNAs. All significance levels in Figure 3A, B and C are based on Wilcox test.

## Data Availability

All scripts and data are available online at https://github.com/gao-lab/RiboCalc/.

## Acknowledgments

The authors thank Drs. Zemin Zhang, Cheng Li, Letian Tao, Jian Lu and Liping Wei at Peking University for their helpful comments and suggestions during the study. Part of the analysis was performed on the Computing Platform of the Center for Life Sciences of Peking University and supported by the High-performance Computing Platform of Peking University.

## Key Points

- We built an *in silico* model for predicting transcripts’ coding ability accurately in human.
- We showed, quantitatively, that both intrinsic and contextual features contribute to coding ability determination.
- We identified a great number of transcripts are with distinct coding abilities among different type of cells (i.e. context-dependent coding transcripts, CDCTs), suggesting the transcript’s coding ability should be modeled as a context-dependent continuous spectrum, rather than a static binary classification as “coding” or “noncoding”.

