## Supplementary Figure 1-12, Supplementary Table 1-8 for "Quantitative model suggests both intrinsic and contextual features contribute to the transcript coding ability determination in cells"

### Supplementary Figures

#### Supplementary Figure 1

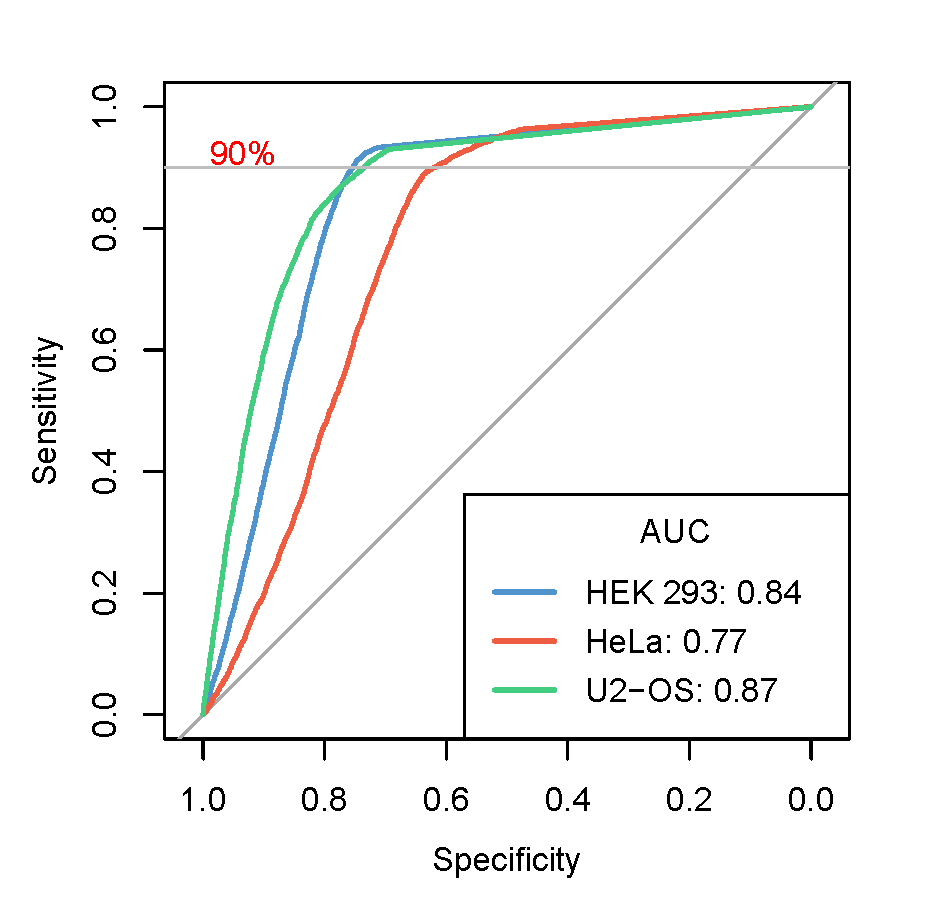

**The ROC curves of the ribORF output p-value with the MS results.** Taken MS identified coding transcripts as positive calling, we evaluated the performance of Ribo-seq based calling tool ribORF (see “**Mass spectrometry (MS) data analysis**” in “**Materials and Methods**”). The X axis shows 1-specificity, and the y axis shows sensitivity. The blue curve was calculated from sample SRR1630831 of HEK 293 cells. The red curve was from SRR970588 of HeLa cells. The green curve was from SRR1551155 of U2-OS cells. The gray horizontal line shows a sensitivity (recall) of 90%. The area under the curve (AUC) of the 3 cell lines is shown in the bottom-right box. The ribORF p-value 0.5 cutoff results in 77%, 72%, 75% specificity with 90%, 90%, 89% sensitivity in HEK 293, HeLa and U2-OS cells, respectively.

#### Supplementary Figure 2

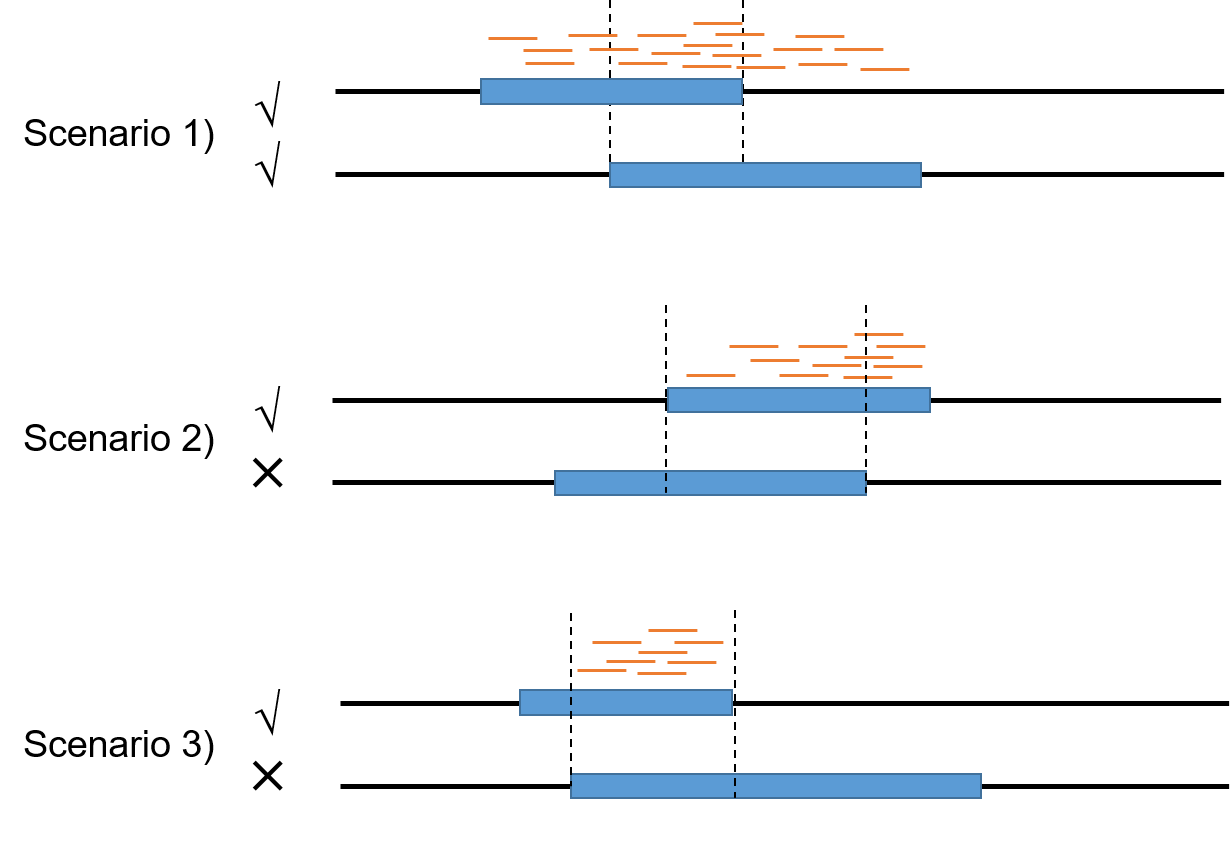

**Most like translated ORF selection procedure for overlapping ORFs**. Scenario 1) overlapping translated ORFs with unique regions covered by ribo-reads were both retained. Scenario 2) the ORFs without unique ribo-reads covering region were filtered out when their overlapping ORFs had. Scenario 3) if the unique regions of both overlapping ORFs were not covered by ribo-reads, the shorter ORF was left.

#### Supplementary Figure 3

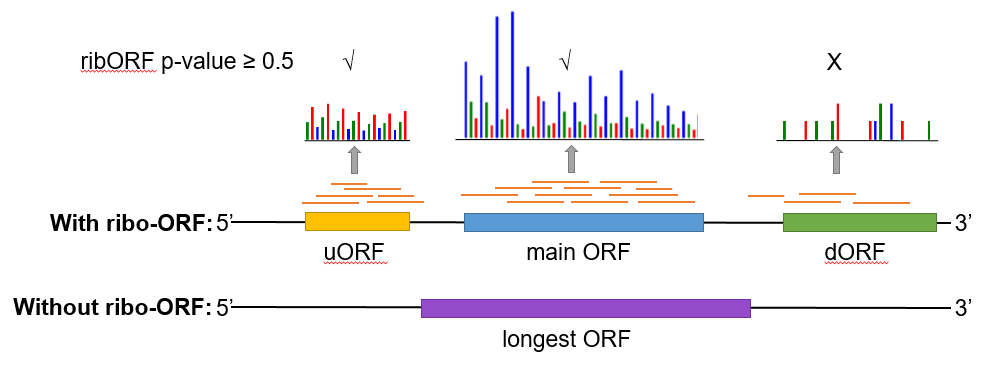

**Translated ORF identification strategies.** We identified translated ORFs as ORFs supported by the Ribo-seq based method, i.e., ribo-ORFs, regardless of whether it was the main ORF annotated based on current knowledge. Thus, it could be an upstream ORF (uORF) or downstream ORF (dORF) of the main ORF. For transcripts without ribo-reads or Ribo-seq data, we took the longest ORF as the putative translated ORF for further calculation.

#### **Supplementary Figure 4**

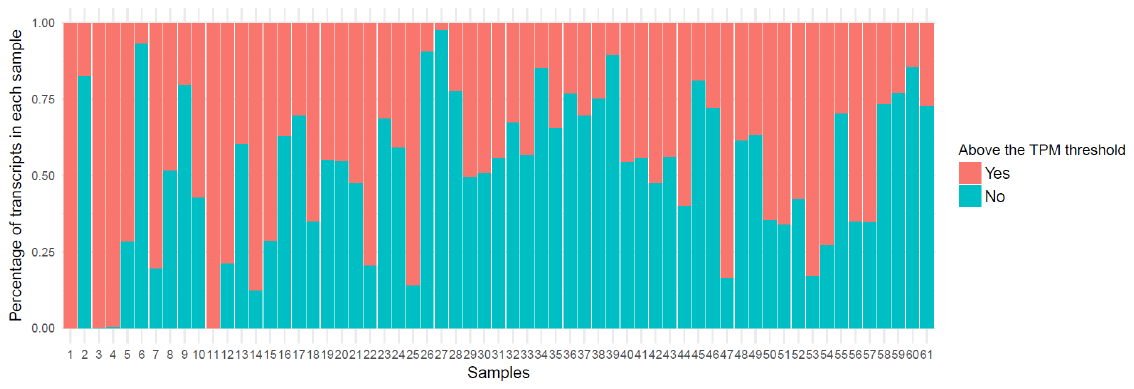

**Fraction of removed noncoding transcripts below the TPM cutoff.** The green fraction was removed noncoding transcripts with a TPM above 0 but below the cutoff of each sample. The red fraction shows the remaining noncoding transcripts. The TPM thresholds for each sample are in **Supplementary Table** **1**.

#### Supplementary Figure 5

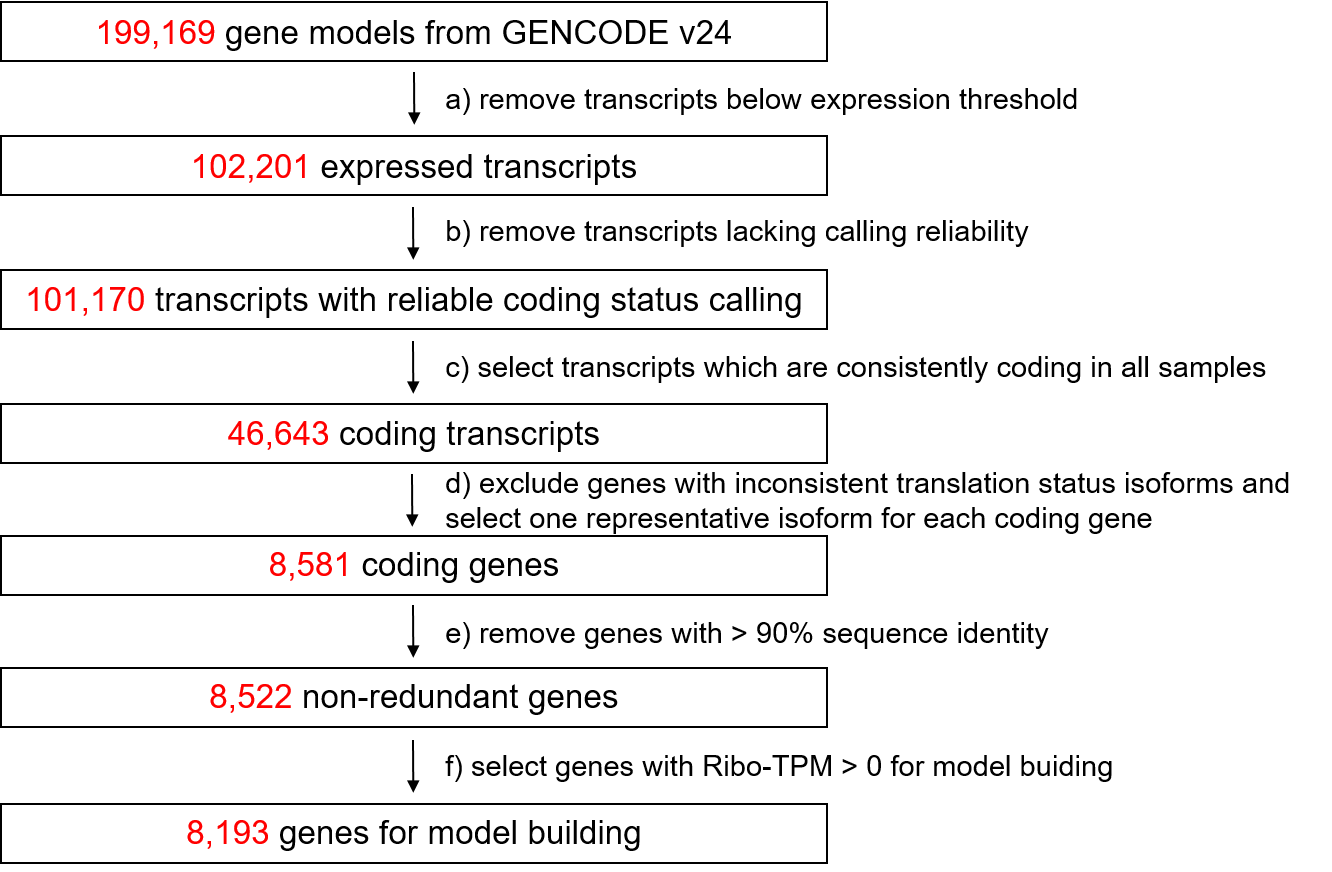

**Workflow of transcripts classification and model data collection.** The red numbers in the boxes show the count of quires left. We collected *bona fide* coding genes for model building using rigorous criteria as followed. **a)** Expression thresholds were used to remove genes with extremely low expression which are not sufficient enough for ribosome capturing. **b)** For calling reliability, we required translated ORF covered by ribo reads with 3-nt periodicity which indicates ribosome move through codons. Thus, 1031 transcripts covered by ribo reads but no 3-nt periodicity were excluded, because we are not able to confirm whether they are translating or just randomly bind with ribosomes. **c)** We only left transcripts consistently coding in all expressed samples. **d)** We left genes that all expressed isoforms were “coding” and selected the isoform with the broadest expression as a representative query for coding gene, because isoforms from same genes often share high sequence similarity. **e)** We removed transcripts with more than 90% sequence using CD-hit. **f)** Finally, we removed coding genes that Ribo-TPM estimated as zero by stringtie. As a result, only 8,193 genes were left for RiboCalc building. For cell-specific models, we only used genes expressed in the corresponding sample from the 8,193 quires which resulting in a lower number of genes.

#### Supplementary Figure 6

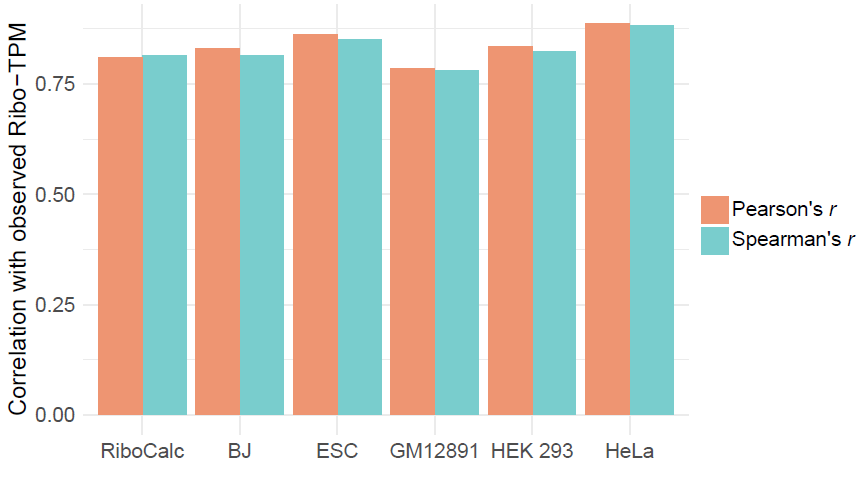

**Performance comparison between RiboCalc and cell-specific models.** The bars show the Pearson’s correlation and Spearman’s correlation between the predicted Ribo-TPM and observed value in cell-specific models and RiboCalc. The transcript data for this plot consist of the testing data in Table 1.

#### Supplementary Figure 7

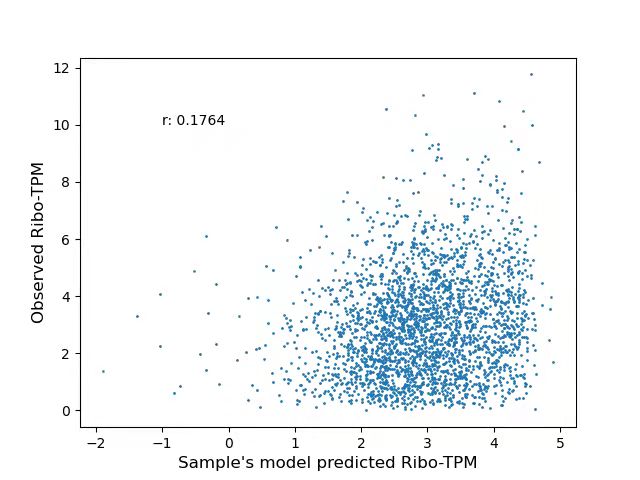

**Predicted Sample *et al.*’s model predicted value on RiboCalc testing data.** By applying Sample’s model directly on RiboCalc testing data (see “**Human model comparison**” in “**Materials and Methods**”), it showed a much lower correlation than RiboCalc (*r* = 0.81).

#### Supplementary Figure 8

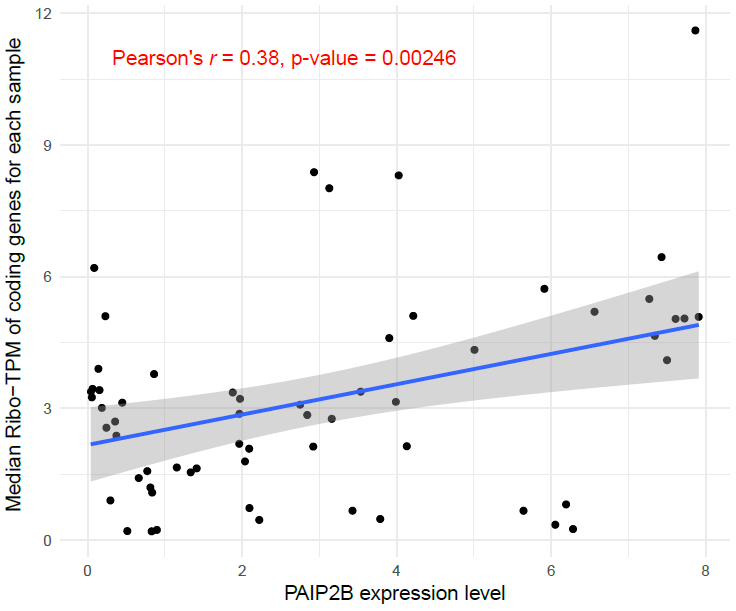

**The expression level of *PAIP2B* and the median Ribo-TPM in each sample.** The x axis shows the RNA TPM of the *PAIP2B* gene in each sample. The y axis shows the median Ribo-TPM of coding genes in each sample. The correlation and its significance level are shown in red text. The blue line is a regression line of x and y fitted via OLS.

#### Supplementary Figure 9

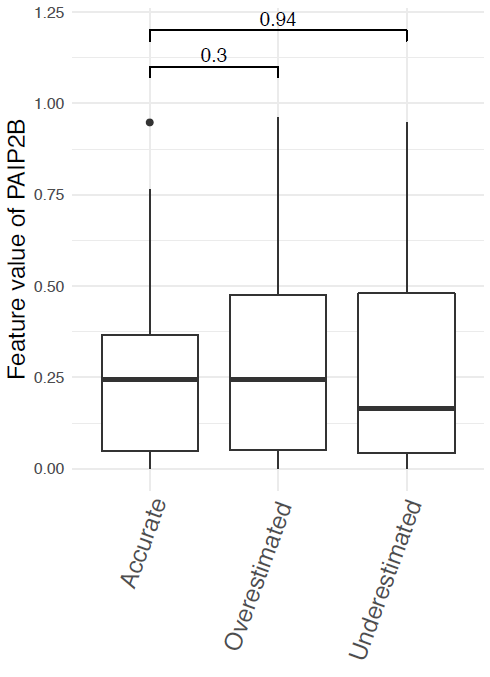

**Feature value of** ***PAIP2B* expression among** **transcripts with different residuals.** The boxes show the feature value of *PAIP2B* expression. “Accurate” refers to 100 transcripts with the lowest absolute value of prediction residuals in the testing set. “Overestimated” refers to 100 transcripts with the most negative prediction residuals, while “Underestimated” refers to 100 transcripts with the most positive prediction residuals. In case of inexplicable parameters caused by a biased model performance, we systematically examined the feature value of transcripts with different prediction residuals but found no significant difference in feature values of *PAIP2B* between accurately predicted transcripts and those with larger residuals.

#### Supplementary Figure 10

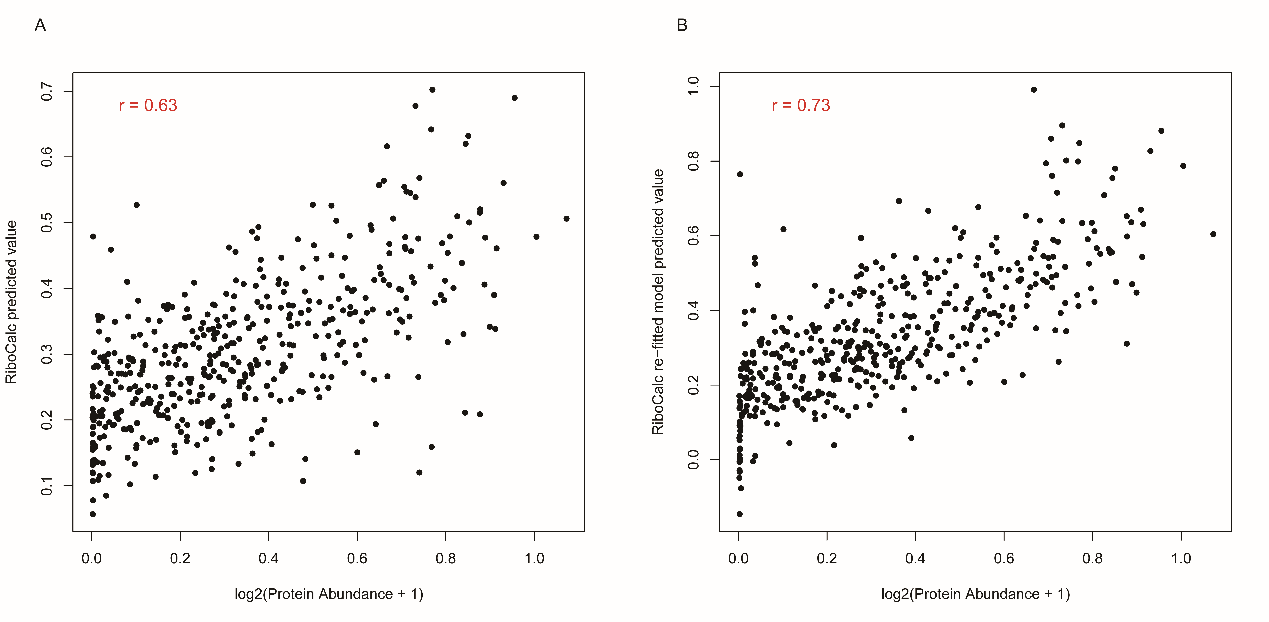

**RiboCalc predicted value and measured protein abundance on testing data.** Protein abundance was downloaded from <https://pax-db.org/dataset/9606/329/> (see “**Testing on OCTOPOS data**” in “**Materials and Methods**”). **A)** Prediction of OCTOPOS data directly using RiboCalc model **B)** Prediction of testing data using model refitted to protein abundance with RiboCalc feature

#### Supplementary Figure 11

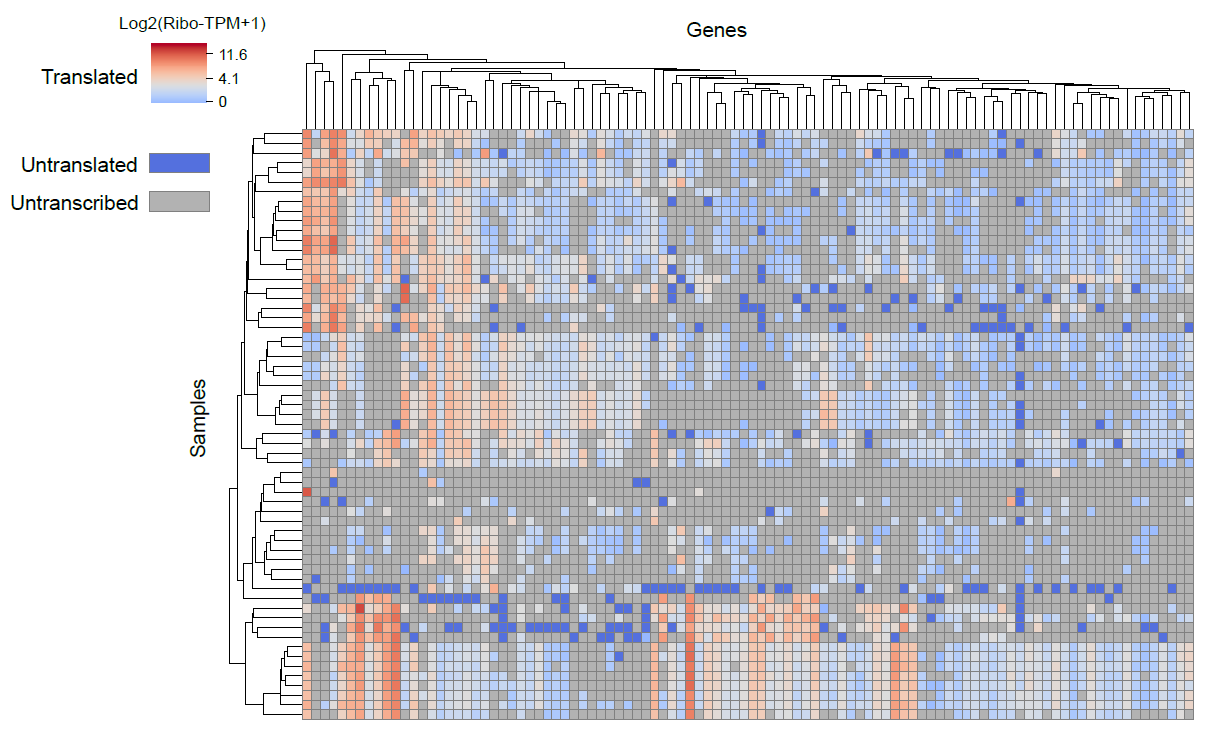

**CDCTs show display various coding ability in different cells.** The x axis shows top 100 CDCT genes expressed in most samples. The y axis shows 61 collected samples. The color in the tile refer to log2 transformed Ribo-TPM. The dark blue tiles show genes transcribed without protein expression (untranslated). The grey tiles show genes that not transcribed.

#### Supplementary Figure 12

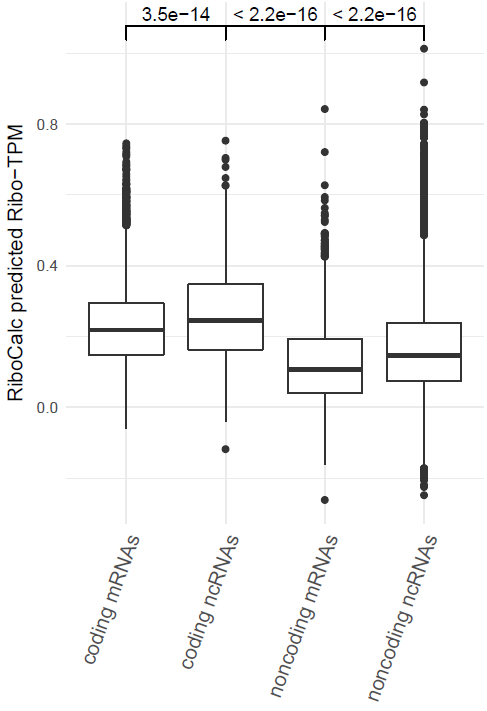

**RiboCalc prediction of transcripts with particular coding ability annotations.** The boxes show the RiboCalc-predicted value with the features of a particular translation-related process. The transcript classifications of “coding mRNAs”, “coding ncRNAs”, “noncoding mRNAs” and “noncoding ncRNAs” are described in Supplementary Table 8. The significance level was based on the Wilcox test. Combining the GENCODE classification and coding ability inferred by RiboCalc, we further group GENCODE transcripts into 4 classes: “coding mRNAs”, “coding ncRNAs”, “noncoding mRNAs” and “noncoding ncRNAs” (Supplementary Table 8), and both coding mRNAs and coding ncRNAs showed a significantly higher RiboCalc scores than of noncoding mRNAs and ncRNAs.

### Supplementary Tables

#### Supplementary Table 1

**Ribo-seq data and the corresponding RNA-seq data used by RiboCalc, and the TPM thresholds used for each sample**

(see Supplementary_Table_1_SampleInformation.xlsx)

#### Supplementary Table 2

**Corresponding Ribo-seq sample for MS data used in comparison**

| **Ribo-seq sample ID** | **MS cell line** |
| --- | --- |
| SRR1630831 | HEK 293 |
| SRR970490 | HeLa |
| SRR970538 |  |
| SRR970561 |  |
| SRR970565 |  |
| SRR970587 |  |
| SRR970588 |  |
| SRR1551155 | U-2 OS |

The HeLa Ribo-seq sample SRR1042900 was removed from the comparison because of adding control siRNAs.

#### Supplementary Table 3

**Recall and precision of Ribo-seq based identification approaches with MS method**

| **MS cell line** | **Ribo-seq sample ID** | **Sensitivity** | | | |
| --- | --- | --- | --- | --- | --- |
|  |  | **RiboCode** | **ribORF≥0** | **ribORF≥0.5** | **ribORF≥0.7** |
| HEK 293 | SRR1630831 | 19.0% | 93.2% | 90.6% | 85.6% |
| U-2 OS | SRR1551155 | 17.8% | 92.8% | 89.0% | 83.4% |
| HeLa | SRR970490 | 13.4% | 95.7% | 87.7% | 65.3% |
|  | SRR970538 | 17.7% | 94.9% | 89.8% | 77.2% |
|  | SRR970561 | 12.7% | 93.4% | 88.3% | 69.8% |
|  | SRR970565 | 26.0% | 95.8% | 90.8% | 86.0% |
|  | SRR970587 | 17.3% | 92.8% | 86.3% | 75.9% |
|  | SRR970588 | 24.9% | 95.3% | 89.5% | 85.2% |
| **MS cell line** | **Ribo-seq sample ID** | **Specificity** | | | |
|  |  | **RiboCode** | **ribORF≥0** | **ribORF≥0.5** | **ribORF≥0.7** |
| HEK 293 | SRR1630831 | 95.6% | 73.3% | 76.6% | 78.3% |
| U-2 OS | SRR1551155 | 95.4% | 70.8% | 75.3% | 80.5% |
| HeLa | SRR970490 | 96.6% | 56.9% | 72.5% | 83.1% |
|  | SRR970538 | 96.0% | 67.7% | 73.3% | 81.3% |
|  | SRR970561 | 96.7% | 63.5% | 70.2% | 78.4% |
|  | SRR970565 | 92.1% | 57.6% | 67.1% | 71.5% |
|  | SRR970587 | 96.0% | 68.1% | 75.3% | 80.2% |
|  | SRR970588 | 93.4% | 60.4% | 71.9% | 74.4% |

The criteria of ribORF≥0, 0.5 or 0.7 refer to ribORF results with output p-value above 0, 0.5 or 0.7 respectively.

#### Supplementary Table 4

**Classification and calculation of candidate features**

| **Related processes** | **Feature name** | **Calculation approach** | **Feature number** |
| --- | --- | --- | --- |
| Expression abundance | RNA TPM | Transformed as log2(TPM+1). The expression abundance (TPM) was estimated by stringtie[1] (see <https://github.com/gao-lab/RiboCalc/blob/master/feature_calculation/RNAandRibo-seq_processing.txt>). | 1 |
|  | AGO CLIP | Binary value of whether the transcripts exon regions covered by AGO1 or AGO2 supported by CLIP-seq data. The CLIP data collection and analysis were implemented by AnnoLnc[2, 3] (see <http://annolnc.gao-lab.org/methods.php#link-anno-pi> for details). | 1 |
|  | miRNA target | Targeting number with human mature miRNAs in 3’UTR region. The target sites were predicted according to miRBase[4] by TargetScan[5]. TargetScan context+ score below -0.2 was taken as cutoff. Overlapped miRNA target region were taken as one target. | 1 |
| Translation initiation | Initiation folding energy | Minimum free energy of upstream and downstream 100nt RNA sequences around start codon predicted by RNAfold[6] | 1 |
|  | Translation initiation motif | Binary value of the existence of translation initiation motif with particular stat codon. fimo[7]. The motifs were identified by TITER [8]. | 4 |
| Translation elongation | Condon frequency | The percentage of particular codon used in the translated ORF | 64 |
|  | CAI | Codon Adaptation Index[9] calculated with EMBOSS package[10] | 1 |
|  | ENC | Effective Number of Codons[11] calculated with EMBOSS package | 1 |
|  | MTDR | Mean of the Typical Decoding Rates [12] calculated by MTDRcalculator[12] with the parameter of human HEK 293 cell line. MTDRcalculator was downloaded from <https://www.cs.tau.ac.il/~tamirtul/MTDR/>. | 1 |
| Translation regulators | Translation-related factor abundance | The RNA TPM of genes with molecular function as “translation regulator activity” (GO:0045182) in GO. The genes not expressing in all the samples were excluded. | 140 |
| Transcript structure | Transcript length | Length of full transcript with log2 transformation | 1 |
|  | CDS length | Transformed as log2(CDS+1). CDS refers to length of translated ORFs | 1 |
|  | 5’UTR length | Transformed as log2(5’UTR+1). 5’UTR refers to length of 5’UTR | 1 |
|  | 3’UTR length | Transformed as log2(3’UTR+1). 3’UTR refers to length of 3’UTR | 1 |
|  | 5’UTR GC | GC content in 5’UTR sequence | 1 |
|  | 3’UTR GC | GC content in 3’UTR sequence | 1 |

The detailed feature calculation scripts are at <https://github.com/gao-lab/RiboCalc/tree/master/feature_calculation>.

#### Supplementary Table 5

**Differences of feature calculation with yeast and human model**

| **Feature name** | **Calculation approach in yeast** |
| --- | --- |
| RNA TPM | Directly retrieved from ‘nar-00812-a-2017-File019.csv’ file in Li *et al.*’s supplementary data. |
| Translation initiation motif | Not used in yeast, because the motifs were specifically identified in human. |
| AGO CLIP | Not used for no CLIP evidences for AGO binding in yeast. |
| miRNA targeting | Not used for no comprehensive annotation for miRNA targeting in yeast. |
| CAI | Calculated by EMBOSS package with the parameter of “-cfile Eyeast.cut”. |
| MTDR | Calculated by MTDRcalculator with the parameter of “S. cerevisiae Exp”. |
| Translation-related factor abundance | Not used in yeast for being useless in unicellular organisms. |
| CDS length | The CDS sequences were retrieved from ‘nar-00812-a-2017-File019.csv’ file in Li *et al.*’s supplementary data. |
| 5’UTR length, 5’UTR GC | The 5’UTR sequences were retrieved from ‘nar-00812-a-2017-File019.csv’ file in Li *et al.*’s supplementary data. |
| 3’UTR length, 3’UTR GC | The 3’UTR sequences were downloaded from S288C strain in SGD database (<https://downloads.yeastgenome.org/sequence/S288C_reference/SGD_all_ORFs_3prime_UTRs.fsa>). When more than one 3’UTR sequences were annotated, we took the longest one. |
| Transcript length | The total length of 5’UTR, CDS and 3’UTR. |

#### Supplementary Table 6

**Correlation between predicted and observed values of RiboCalc and Li’s human model**

| **Predicted value** | **Model** | **Pearson’s *r*** | **Spearman’s *r*** |
| --- | --- | --- | --- |
| TR | RiboCalc | 0.659 | 0.683 |
|  | Li human | 0.644 | 0.667 |
| Ribo-RPKM | RiboCalc | 0.923 | 0.925 |
|  | Li human | 0.921 | 0.922 |

#### Supplementary Table 7

**Feature coefficients of RiboCalc human, yeast and cell-specific models**

(see Supplementary_Table_7_FeatureCoeficient.xlsx)

1. The feature names are listed in the following order: translation initiation, translation elongation, expression abundance, transcript structure, translation regulators.
2. The “.” in the table refer to no these features in the corresponding mode. For instance, features of translation regulators are unnecessary for cell-specific model.
3. The “removed” in the table refer to the features were removed for the existence of highly correlated features.

#### Supplementary Table 8

**Coding ability classification for transcripts based on GENCODE and ribosome profiling data**

|  | **Coding in GENCODE** | **Noncoding in GENCODE** |
| --- | --- | --- |
| **Coding in Ribo-seq** | coding mRNAs | coding ncRNAs |
| **Noncoding in Ribo-seq** | noncoding mRNAs | noncoding ncRNAs |
